# Spontaneous Mutants of *Streptococcus sanguinis* with Defects in the Glucose-PTS Show Enhanced Post-Exponential Phase Fitness

**DOI:** 10.1101/2021.07.15.452590

**Authors:** Lin Zeng, Alejandro R. Walker, Kyulim Lee, Zachary A. Taylor, Robert A. Burne

**Author notes:** Correspondence Lin Zeng, PhD.

## Abstract

Genetic truncations in a gene encoding a putative glucose-PTS protein (*manL*, EIIAB^Man^) were identified in subpopulations of two separate laboratory stocks of *Streptococcus sanguinis* SK36; the mutants had reduced PTS activities on glucose and other monosaccharides. Using an engineered mutant of *manL* and its complemented derivative, we showed that the ManL-deficient strain had improved bacterial viability in stationary phase and was better able to inhibit the growth of the dental caries pathogen *Streptococcus mutans*. Transcriptional analysis and biochemical assays suggested that the *manL* mutant underwent reprograming of central carbon metabolism that directed pyruvate away from production of lactate, increasing production of hydrogen peroxide (H_2_O_2_) and excretion of pyruvate. Addition of pyruvate to the medium enhanced the survival of SK36 in overnight cultures. Meanwhile, elevated pyruvate levels were detected in the cultures of a small, but significant percentage (∼10%), of clinical isolates of oral commensal bacteria. Furthermore, the *manL* mutant showed higher expression of the arginine deiminase system than the wild type, which enhanced the ability of the mutant to raise environmental pH when arginine was present. Significant discrepancies in genome sequence were identified between strain SK36 obtained from ATCC and the sequence deposited in GenBank. As the conditions that are likely associated with the emergence of spontaneous *manL* mutations, i.e. excess carbohydrates and low pH, are those associated with caries development, we propose that the glucose-PTS strongly influences commensal-pathogen interactions by altering the production of ammonia, pyruvate, and H_2_O_2_.

**Importance:** A health-associated dental microbiome provides a potent defense against pathogens and diseases. *Streptococcus sanguinis* is an abundant member of a health-associated oral flora that antagonizes pathogens by producing hydrogen peroxide. There is a need for a better understanding of the mechanisms that allow bacteria to survive carbohydrate-rich and acidic environments associated with the development of dental caries. We report the isolation and characterization of spontaneous mutants of *S. sanguinis* with impairment in glucose transport. The resultant reprograming of central metabolism in these mutants reduced the production of lactic acid and increased pyruvate accumulation; the latter enables these bacteria to better cope with hydrogen peroxide and low pH. The implications of these discoveries in the development of dental caries are discussed.

## Introduction

Dental caries is caused by dysbiosis in the dental microbiome, where an overabundance of acid-producing (acidogenic) and acid-resistant (aciduric) bacteria such as mutans streptococci and lactobacilli, along with certain *Actinomyces, Scardovia*, and fungal species, drives the acidification of dental biofilms and demineralization of tooth enamel. Diets rich in carbohydrates are critical to caries formation, while host genetics and socioeconomic factors also affect the incidence and severity of the disease(s) (1, 2). Organic acids, including lactic, acetic, and formic, are some of the primary products released by oral bacteria that ferment carbohydrates, which include bacteria that are considered etiological agents of caries and those that are considered commensals. Due to its low p*Ka* value, lactic acid is particularly damaging to tooth enamel. When dietary carbohydrates are ingested, lactate can be produced in large quantities by a group of oral streptococci that includes *Streptococcus mutans*, the primary etiological agent of dental caries. Other bacterial factors important to the ecological balance of the dental microbiome include reactive oxygen species, e.g. hydrogen peroxide (H_2_O_2_) and alkaline compounds, such as ammonia. Many bacteria, *S. mutans* in particular, are sensitive to physiologically relevant concentrations of H_2_O_2_, and thus can be inhibited by the presence of peroxigenic commensals, mainly mitis group of streptococci, which includes *S. sanguinis, Streptococcus gordonii, Streptococcus mitis* and other commensal streptococci that are among the most abundant members of the dental microbiome (3). Generally, *S. sanguinis* and *S. gordonii* are considerably less acid tolerant (aciduric) than *S. mutans*, but carry the arginine deiminase (AD) system, which in the presence of arginine, releases ammonia and provides ATP to improve the survival and persistence of these organisms when faced with an acid challenge (4). Past research has indicated that production of both H_2_O_2_ (5–7) and AD activities (8) by these commensals can be influenced by bacterial uptake and catabolism of specific carbohydrates.

For most oral streptococci, carbohydrates are primarily internalized via the phosphoenolpyruvate::sugar phosphotransferase system (PTS), which is composed of two general proteins, Enzyme I (EI) and the phospho-carrier protein HPr, and a variety of carbohydrate-specific Enzymes II (EII) that are membrane-associated permeases (9). The PTS concurrently internalizes and phosphorylates carbohydrates that can be fed into the Embden-Meyerhoff-Parnas (EMP) pathway, which primarily yields pyruvate and the energy molecules ATP and NADH. To maintain bacterial redox balance and resupply glycolysis, NADH must be oxidized back into NAD^+^. In oral streptococci, this can occur via lactate dehydrogenase (LDH) (10), an NADH oxidase (NOX), or other redox-coupled reactions (11). As the central point for bacterial energy metabolism and biogenesis, pyruvate can supply the tricarboxylic acid cycle (TCA) for bacteria that can conduct aerobic respiration, or alternatively can be converted to the aforementioned organic acids when oxygen is limited. As most of the streptococci have only a partial TCA cycle and lack cytochromes, the fate of pyruvate is limited to either homolactic fermentation, yielding lactic acid via the reducing activity of LDH, or heterofermentation that can produce ethanol, acetic acid, formate, and other end products depending on conditions and carbohydrate source(s)(12, 13). Oxidation of pyruvate by a few non-LDH pathways, including pyruvate dehydrogenase (PDH), pyruvate-formate lyase (Pfl), and pyruvate oxidase (POX) allows the bacteria to create more energy molecules and produce end products with either milder or no acidic properties. It is primarily the activity of POX, encoded by *spxB* in *S. sanguinis*, that produces H_2_O_2_; *S. mutans* lacks *spxB* and does not produce H_2_O_2_ in any significant quantities. Distribution of pyruvate between the LDH and non-LDH pathways is regulated at both the transcriptional and enzymatic levels in response to bacterial energy status, generally metabolic intermediates such as fructose-1,6-bisphosphate (F-1,6-bP), and environmental cues such as carbohydrate abundance, pH, and oxygen levels (10, 12). The LDH pathway is usually favored under low-oxygen and carbohydrate-excess conditions.

As one of the early colonizers of oral cavity, *S. sanguinis* is an abundant commensal species that is frequently associated with oral health (14). It is also considered an opportunistic pathogen of infective endocarditis (IE). Previous research has indicated that *S. sanguinis* can ferment a large array of carbohydrates for acid production (15). *In silico* analyses also identified PTS transporters and metabolic pathways that are comparable to those in other more-intensively studied streptococcal species, such as *S. mutans* (16, 17). Notably different from *S. mutans*, *S. sanguinis* possesses the genes necessary to carry out gluconeogenesis, oxidation of pyruvate to produce H_2_O_2_, and the full functions of the pentose phosphate pathway. Recently, a transcriptomic study delineated in *S. sanguinis* SK36 the regulon governed by the transcriptional regulator CcpA (18), which in many Gram-positive Firmicutes is the dominant regulator of carbohydrate catabolite repression (CCR), a phenomenon where the genes encoding metabolic pathways for non-preferred carbohydrates are transcriptionally suppressed until a preferred carbohydrate such as glucose has been exhausted (19). Loss of CcpA in *S. sanguinis* SK36 affected expression of nearly 20% of the genome. Interestingly, studies on the effects of *ccpA* deletion in *S. sanguinis* reported enhanced secretion of H_2_O_2_ without a significant change in antagonistic potential against *S. mutans*. Expression of *spxB* was shown to be regulated by CcpA, but the increased secretion by the *ccpA* mutant of pyruvate, an antioxidant, was postulated to counteract the impact of overproduction of H_2_O_2_ (20, 21).

We recently identified a memory effect of sugar metabolism in *S. mutans*, where past catabolism of monosaccharides, such as glucose and fructose, had profound impacts on the capacity of the bacterium to utilize lactose (22). To ascertain whether the memory effect was a general behavior of oral streptococci, these investigations were expanded to include *S. sanguinis* SK36, and a deletion mutant of the lactose repressor LacR was constructed to study its function in regulating catabolism of multiple carbohydrates. Here, we report the unexpected identification of spontaneous glucose-PTS mutations in two SK36 stocks that afforded a subpopulation of the cells enhanced fitness under laboratory conditions. Further analysis indicated that the PTS plays an important role in the regulation of central carbon metabolism in *S. sanguinis* in ways that contribute to the ability of this commensal to persist under stress and to compete against the pathobiont *S. mutans*.

## Results

### Isolation from SK36 of spontaneous mutants deficient in glucose PTS

We previously investigated the impact on carbohydrate metabolism of deletion of a lactose repressor gene *lacR* in *S. sanguinis* SK36 (22). When analyzing growth and carbohydrate transport in the *lacR* mutant of SK36, we noted an unusually severe defect in PTS activity compared to similarly constructed *lacR* mutants we had created in related bacteria. In an effort to understand the basis of this phenotype, we conducted whole genome sequencing (WGS) of the *lacR* mutant and, as a control, the parental strain of frozen stock of SK36 (here designated as strain MMZ1612). Results of the WGS indicated the presence in our wild-type laboratory strain of SK36 (MMZ1612) of 115 single-nucleotide polymorphisms (SNPs, Table S1) that were not present in the published genome of SK36 available at GenBank. Notably, there were two truncations, one being a 330-bp deletion in a putative glucose-PTS gene (SSA_1918, tentatively identified as *manL*) that encodes the A and B domains of the Enzyme II of a PTS permease (23), and the other a 342-bp deletion in an open reading frame (ORF) (SSA_1927) that encodes for a putative transporter predicted to confer tellurite resistance (24). The SSA_1918 would have resulted in a translational product that terminates before the EIIB domain of the apparent ManL homologue. Two PCR reactions were designed based on this information and used to assess the integrity of the *manL* gene and SSA_1927 in random isolates selected from individual colonies grown from our MMZ1612 frozen stock. The results provided evidence that these two deletions were likely present in a subpopulation of the stock, comprising roughly 30% of the viable cells.

Subsequently, a second attempt was made at creating the *lacR* deletion, by requesting a different stock of SK36 (here designated MMZ1896) from the laboratory of Todd Kitten, which works extensively with SK36. After performing the same genetic manipulation, similar growth phenotypes (data not shown) were noted in some of the *lacR*-null clones that again suggested a deficiency in glucose-PTS activity. WGS, followed by PCR for selected regions of the genome, indeed identified a nonsense mutation event, Q217* (CAA→TAA), in ∼10% of the population that resulted in truncation of the SSA_1918/*manL* gene at approximately the same location as in MMZ1612. Similar to our SK36 stock (MMZ1612), 114 SNPs were identified in MMZ1896 (Kitten stock; Table S1), which, except for one, matched what was found in our freezer stock (MMZ1612). The genetic lesions in *manL* could reduce the capacity of *S. sanguinis* to utilize a number of carbohydrates commonly present in the human oral cavity, since the ManLMN permease in *S. mutans* is the primary transporter for glucose, galactose, glucosamine (GlcN) and N-acetylglucosamine (GlcNAc) (see Fig. S1 for growth curves). Indeed, when a wild-type copy of the *lacR* gene was later introduced into one such *lacR*-null strain using an integration vector pMJB8 (25), the complemented strain remained deficient in growth on glucose and certain other monosaccharides dependent on the ManLMN glucose-PTS for internalization (data not shown). Nonetheless, the fact that these mutants are so well-represented in populations could indicate that there are physiological benefits associated either with the specific lesions in the glucose-PTS or with other not-yet-investigated SNPs that, under certain conditions, outweighed the reduced fitness associated with loss of a primary hexose internalization system.

### Enhanced persistence of *manL* mutants under acidic stress

To further investigate the involvement of the glucose-PTS in the observed behaviors, a number of *manL* deletion mutants were constructed in the backgrounds of wild-type isolates of SK36 (MMZ1612 and MMZ1896) via the allelic exchange method (Table 1). To rule out the possibility of additional spontaneous mutations obscuring the effects of *manL* deletion, two *manL*-complemented derivatives (*manLComp*) were constructed in the *manL-*null background via a “knock-in” approach (26). The wild-type parent SK36, a *manL* mutant and a *manLComp* strain were first studied for their growth phenotypes. Strains were cultivated to exponential phase in BHI before diluting into the chemically-defined medium FMC formulated with glucose, galactose, GlcN or GlcNAc as the sole carbohydrate source. Compared to the wild type and the complemented strain, *manL* mutant strains showed reduced growth rates on glucose and galactose, and especially poor growth on the amino sugars GlcN and GlcNAc (Fig. S2). The growth defects in the *manL* mutant were similar to those of the aforementioned *lacR* mutant (Fig. S1). To assess long term viability and effects of pH thereon, we streaked the strains on BHI agar or BHI agar supplemented with 50 mM potassium phosphate buffer (pH 7.2). Wild-type SK36 remained viable for at least 3 weeks at 4°C on the buffered plates, whereas viable cells could not be recovered from unbuffered plates after a week or less. Conversely, the *manL* mutant survived one to two weeks longer than SK36 on unbuffered BHI agar. When tested for growth characteristics by diluting directly from overnight cultures, deletion of *manL*, or addition of phosphate buffer in the overnight cultures, significantly shortened the lag phase following sub-culture into fresh BHI or into TY medium supplemented with glucose (TY-Glc) (Fig. 1AB). These effects indicated improved viability in the overnight cultures as a result of *manL* deletion, and that loss of viability likely involved exposure to lower pH values than in buffered media. To further examine the relative fitness of the strains, overnight cultures of SK36 and the *manL* mutant were used in a competition assay by mixing in a 1:1 ratio (based on OD_600_) and diluting 1000-fold into fresh BHI medium. After one more round of dilution (at 5 h) followed by an overnight incubation (24 h), cells were plated for CFU enumeration on selective agar plates. Competition indices (CI) were calculated using CFUs at 0 and 24 h time points. Out of the three replicates, *manL* outcompeted the wild-type parent by a large margin in two biological replicates, with CI = 25.1 and 9.9; and in the third replicate SK36 was no longer detected. These results strongly supported the notion of enhanced fitness due to deletion of *manL*, providing a potential explanation for the emergence of glucose PTS-negative isolates under laboratory conditions.

**Figure 1.**
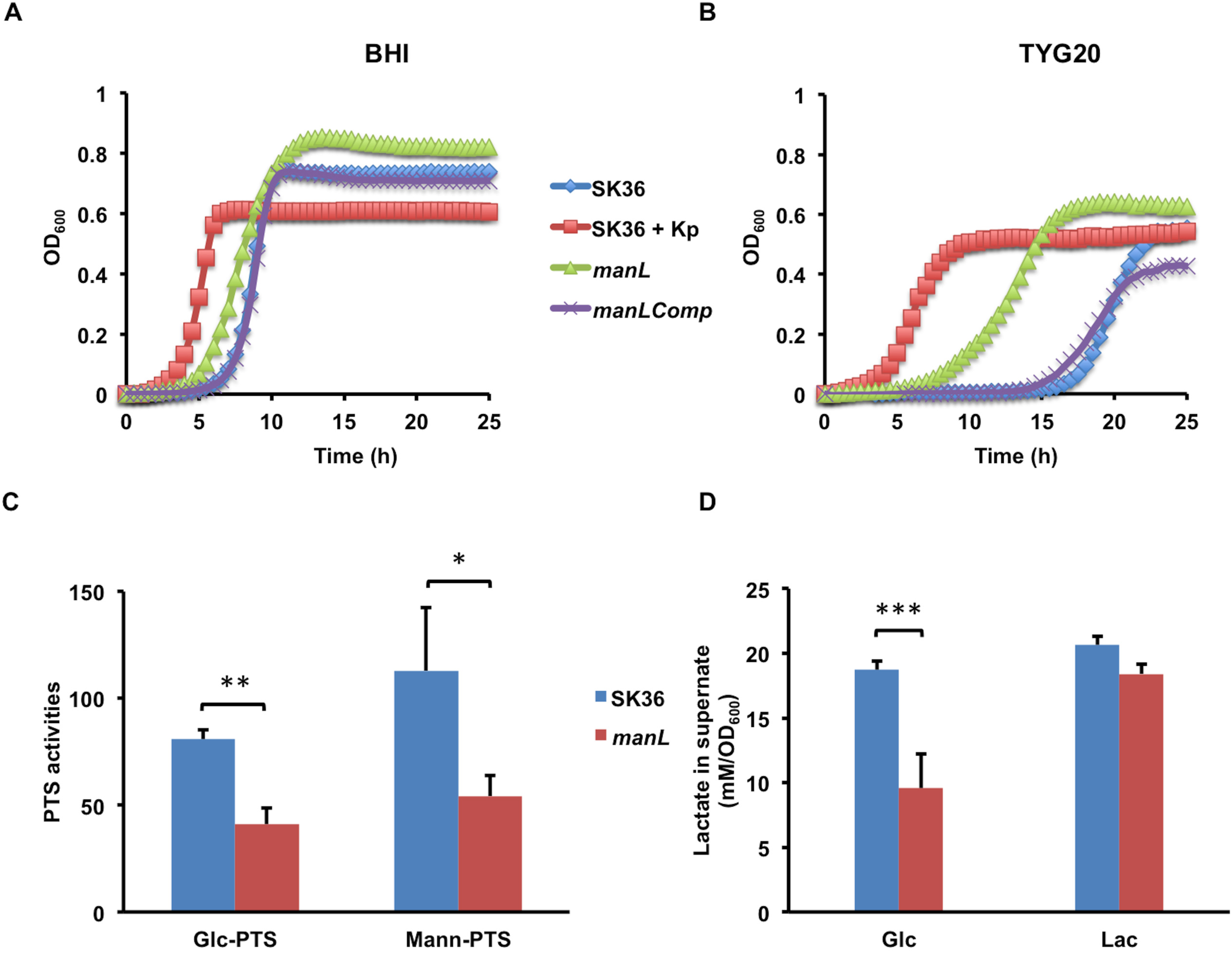
A *manL* mutant of *S. sanguinis* SK36 had enhanced viability in stationary cultures. Wild-type strain SK36 (MMZ1612), the *manL* mutant, and the complemented derivative of *manL* (*manLComp*) were (n = 3) cultivated for 20 h in BHI before being diluted into fresh BHI (A) or a TY medium containing 20 mM of glucose (TYG20) (B), followed by growth monitoring in a Bioscreen system. For another set of SK36 samples (SK36 + Kp), 50 mM of potassium phosphate buffer (pH 7.2) was added to the overnight cultures. (C) An *in vitro* sugar phosphorylation assay was carried out using SK36 and *manL* mutant cultures prepared with BHI medium and harvested at the exponential phase. (D) For measurements of lactate, overnight cultures were diluted into TY containing glucose (Glc) or lactose (Lac), and the supernates were harvested at the exponential phase. The results are the averages of three biological replicates with error bars denoting standard deviations. Asterisks represent statistical significance according to a Student’s *t* test (*, *P* <0.05; **, *P* <0.01, ***, *P* <0.001).

**Table 1.** Bacterial strains used in this study.

| Strains | Relevant characteristics<sup>a</sup> | Source or reference |
| --- | --- | --- |
| MMZ1612 | *S. sanguinis* SK36 stock 1 | Lemos laboratory |
| MMZ1896 | *S. sanguinis* SK36 stock 2 | Kitten laboratory |
| MMZ1922 | *S. sanguinis* SK36 stock 3 | ATCC BAA-1455 |
| MMZ1616 | SK36 *manL::Em* | MMZ1612 |
| MMZ1617 | SK36 *manL::Km* | MMZ1612 |
| MMZ1882 | SK36 *manL::Km* | MMZ1896 |
| MMZ1923 | SK36 *manL::Km* | MMZ1922 |
| MMZ1904 | SK36 *manLComp::Km* | MMZ1616 |
| MMZ1905 | SK36 *manLComp::Em* | MMZ1617 |
| MMZ1911 | SK36 *arcA::Km* | MMZ1896 |
| MMZ1912 | SK36 *arcA::Em manL::Km* | MMZ1617 |
| MMZ1913 | SK36 *ccpA::Km* | MMZ1896 |
| MMZ1910 | SK36 *ccpA::Em manL::Km* | MMZ1617 |
| MMZ1906 | SK36 *nox::Km* | MMZ1896 |
| MMZ1907 | SK36 *nox::Em manL::Km* | MMZ1617 |
| UA159 | *S. mutans* wild type, *perR*<sup>+</sup> | ATCC 700610 |

<sup>a</sup> *Km* indicates kanamycin resistance; *Em*, erythromycin.

A series of biochemical experiments were carried out to compare the acidogenic and aciduric properties of SK36 and the constructed isogenic *manL* mutant. First, PTS assays (Fig. 1C), which measure *in vitro* sugar phosphorylation by permeabilized bacterial cells, showed a significant reduction in the ability of the *manL* mutant to transport glucose or mannose, thus confirming the predicted function of the glucose-PTS operon in which the *manL* gene resides. Next, a pH drop experiment was performed using late-exponential phase cultures prepared with BHI. When provided with 50 mM glucose, the *manL* mutant showed a slightly slower rate of lowering the pH, but the final pH attained by the mutant was slightly lower, by about 0.14 pH units at the 40-min mark, than the wild type (Fig. 2A). The *manLComp* strain produced a resting pH comparable to the *manL* mutant, suggesting that the difference among these strains may not be biologically significant. However, when the same cultures were first frozen at - 80°C, thawed and then their optical density was normalized before the pH drop assay, the *manL* mutant showed a much greater capacity to lower the environmental pH than the wild type, dropping it at a faster rate and producing a significantly lower resting pH at the end of a 1-hour assay (Fig. 2B). Similar results were obtained when cells harvested from TY-Glc cultures were used in pH drop (data not shown). These results suggested that the *manL* mutant was better at maintaining viability and/or metabolic activity after a freeze/thaw cycle.

**Figure 2.**
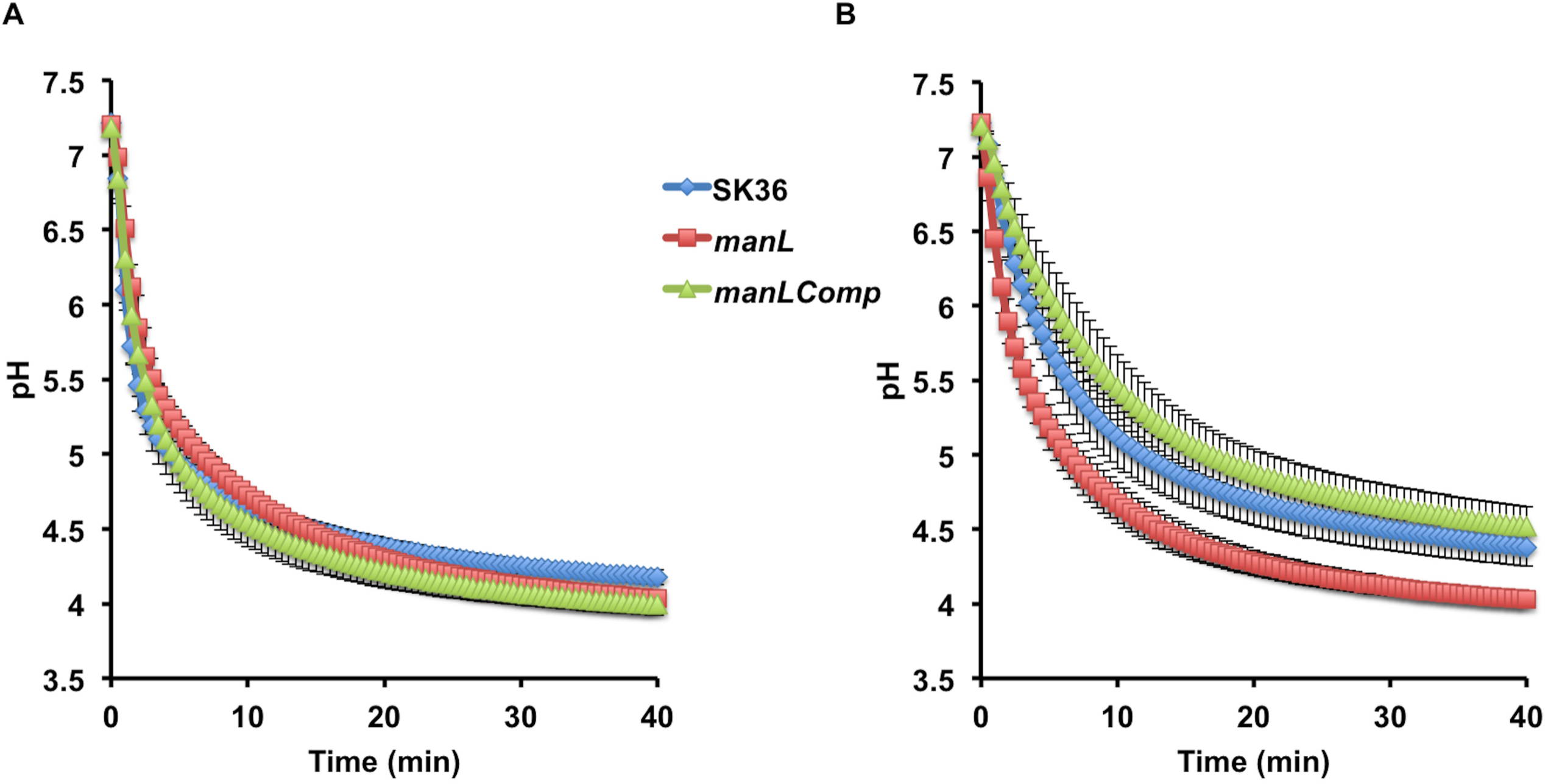
pH drop assays. Strains SK36 (MMZ1612), *manL,* and *manLComp* were cultured to late exponential phase (OD_600_ = 0.8) in 50 mL BHI medium, harvested by centrifugation and used immediately for assay (A) or frozen at - 80°C, and then thawed and assayed at least one day later (B). Each sample was washed once with 50 mL cold water, resuspended in a solution containing 50 mL KCl and 1 mM MgCl_2_, and normalized to OD_600_ = 4.5. The assay was initiated by the addition of 50 mM glucose and pH was monitored and recorded at 30-s intervals for at least 40 min. Each curve includes the average and standard deviation (error bars) of three biological replicates.

Furthermore, when SK36 and the *manL* mutant were assessed for aciduricity by growing on acidified agar plates or acidified BHI broth (adjusted to pH 6.0 and pH 5.5, respectively), there was little to no difference in growth phenotypes between the mutant and the wild type under either condition (data not shown). However, when these strains were subjected to acid killing by incubating in 0.1 M glycine, pH 3.8, the wild type rapidly aggregated, whereas there was no obvious evidence of aggregation in the suspension of the *manL* mutant throughout the 1-h period. Since aggregation could greatly reduce the accuracy of CFU enumeration, we did not assess survival at pH 3.8 by plating. However, both the sensitivity to freezing and thawing and the aggregation differences between the wild-type and *manL* deletion strains point to differences in envelope integrity.

Next, the strains were cultured batch-wise in BHI, in which glucose is the primary carbohydrate source, or in TY prepared with glucose or lactose, and the final pH after ≥20 h of incubation were recorded. These pH measurements showed an intriguing pattern. In overnight TY-Glc cultures, the *manL* mutant achieved a significantly higher final pH than the wild type (Fig. 3A), whereas in BHI cultures the *manL* mutant had a slightly lower pH than the wild type (Fig. 3C). The pH measurements of *manLComp* cultures matched those of the wild type. Meanwhile, little difference in pH was seen among the three strains after overnight growth in TY-lactose (Fig. 3A). This medium-specific effect on final pH after ≥20 h of growth could suggest a potential impact of the glucose-PTS on the ability of the strains to produce alkali via the arginine deiminase (AD) system, which releases ammonia that neutralizes acids both inside and outside of the cells (27). This hypothesis was tested first by RT-qPCR measuring the levels of mRNA for *arcA*, encoding the arginine deiminase enzyme, in cells cultured in TY-Glc or BHI. The results (Fig. 3D) indeed showed a significant increase in expression of *arcA* associated with loss of *manL*. Importantly, in overnight TY-Glc cultures, pH measurements of an *arcA manL* double mutant were significantly lower than the mutant deficient in *manL* alone (Fig. 3B). Therefore, enhanced activities of the AD system were in large part responsible for the increased pH seen in TY-Glc cultures of the *manL* mutant. To reconcile this conclusion with the pH measurements from BHI cultures, we posited that the different impacts of *manL* deletion were due to a lack of significant levels of free arginine in BHI. As a simple test of this hypothesis, L-arginine was added to BHI medium at supplemental concentrations of 1 mM and 5 mM before cultivation of these three strains. The results (Fig. 3C) showed an arginine-dependent increase in final pH in all cultures, with the *manL* mutant yielding higher pH than the wild type when 5 mM supplemental arginine was added. Collectively, these results demonstrated significant roles of the glucose-PTS in regulating aciduricity and alkali generation, thereby affecting the competitiveness of the commensal bacterium under acidic conditions, such as those created by the fermentation of large amounts of carbohydrates.

**Figure 3.**
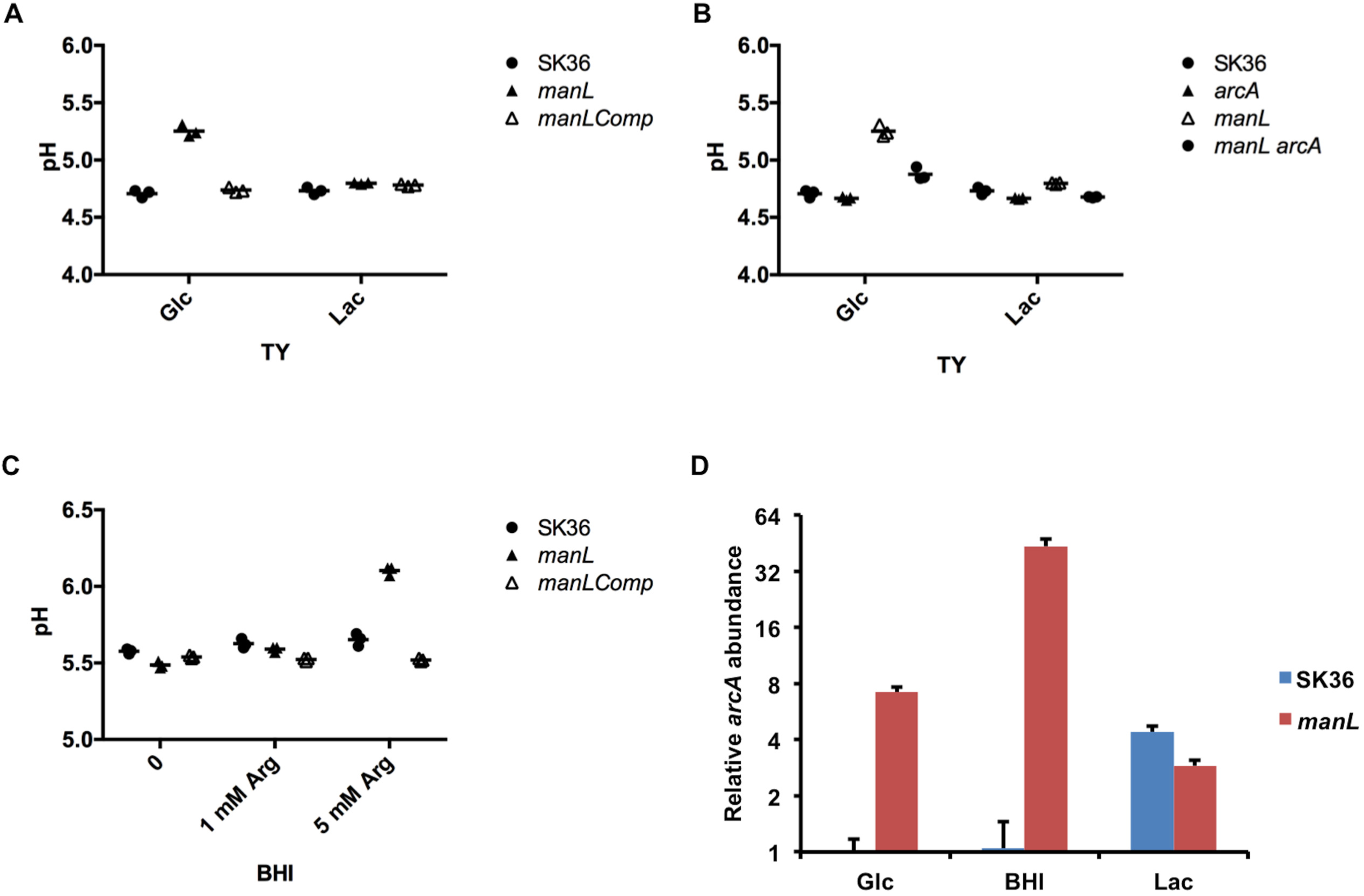
Loss of *manL* affects pH homeostasis by changing AD gene expression. pH measurements of stationary-phase cultures (20 h) in TY (A, B) or BHI (C) media were recorded for strains SK36 (MMZ1612), *manL, manLComp*, *arcA* and *manL arcA.* For BHI cultures, 0, 1, or 5 mM of arginine was added to the medium before cultivation. Data were obtained from three biological replicates. (D) RT-qPCR was performed to measure the expression of *arcA* gene in exponential-phase cultures prepared in TY containing glucose (Glc) or lactose (Lac), or in BHI. The abundance of *arcA* mRNA was calculated relative to an internal control (*gyrA*). Three biological replicates were included for each sample and the results are their averages and standard deviations (as error bars).

### The *manL* mutant produces more H_2_O_2_ and less lactate

*S. sanguinis* is considered a health-associated commensal and has antagonistic properties toward the major etiologic agent of dental caries, *S. mutans*; with a primary antagonistic factor being hydrogen peroxide (H_2_O_2_). When the *manL* deletion mutant and its complemented derivative *manLComp* were tested for their abilities to release H_2_O_2_ on agar plates, visualized as precipitation zones on Prussian blue plates (28), the results showed that loss of *manL* significantly enhanced the release of H_2_O_2_ by the bacterium (Fig. 4A). When tested in a plate-based competition assay together with *S. mutans*, the *manL* mutant showed a significantly increased ability to inhibit *S. mutans* UA159, relative to the wild-type SK36 (Fig. 4B). The UA159 strain used in this study was a *perR*^+^ stock obtained from ATCC, as opposed to the UA159 derivative that carries a spontaneous mutation in *perR* that truncates PerR and reduces sensitivity to H_2_O_2_ and oxidative stress in general (29). When *S. sanguinis* strains were each mixed with UA159 before being placed onto the agar plates, the *manL* mutant similarly outperformed its wild-type parent in competition against UA159, as quantified by CFU enumeration (data not shown). The *manLComp* strain behaved similarly to the *manL*^+^ parent strain in these assays (Fig. S3). Consistent with the role of ManLMN in transporting glucose, both of these phenotypes depended on the presence of glucose (20 mM) and oxygen, and were not seen when lactose (10 mM) was used in place of glucose as the growth carbohydrate (Fig. 4). The *manL* mutant, however, showed enhanced H_2_O_2_ production on TY agar formulated with a combination of glucose and galactose, or with only galactose, GlcN, or GlcNAc (Fig. 4). As noted above, these four carbohydrates are transported primarily via the ManLMN PTS permease in closely related bacteria (30–32).

**Figure 4.**
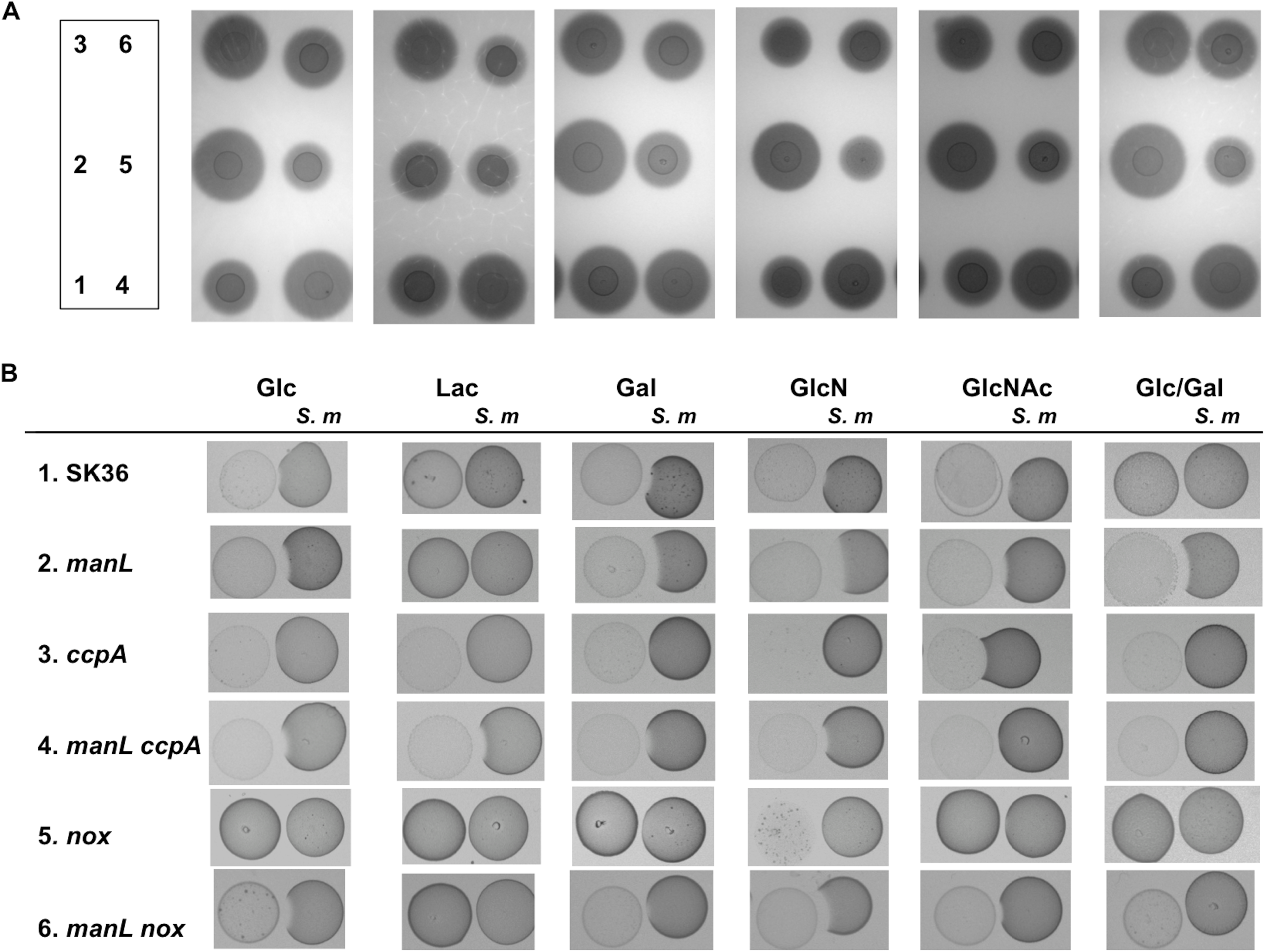
H_2_O_2_ production (A) and antagonism of *S. mutans* (B) on plates. Cultures of SK36 the wild type (strain #1) and its mutant derivatives deficient in *manL* (#2), *ccpA* (#3), *manL ccpA* (#4), *nox* (#5), and *manL nox* (#6) were each dropped onto the surface of TY-agar plates prepared with 20 mM of glucose, lactose, galactose (Gal), GlcN, GlcNAc, or 10 mM each of glucose and galactose (Glc/Gal), and incubated for 24 h in an aerobic environment (with 5% CO_2_). (A) For direct measurement of H_2_O_2_ release, the plates contained 0.1% each of FeCl_3_.6H_2_O and potassium hexacyanoferrate(III) which formed a Prussian blue zone upon reacting with H_2_O_2_. (B) For antagonism of *S. mutans*, same amount of UA159 culture (*S. m*) was placed to the right of the first colony, followed by another 24 h of incubation. All images were photographed under the same setting, with the zones of Prussian blue assessed relative to the size of the colonies. Each experiment was repeated three times using biological replicates, with a representative result being presented.

As H_2_O_2_ is generated by pyruvate oxidase (SpxB) during the conversion of pyruvate to acetyl phosphate (AcP) in the presence of oxygen, we reasoned that deletion of *manL* could have altered the flow of pyruvate in bacterial central metabolism (12). Under carbohydrate-rich and oxygen-limited conditions, streptococci are known to produce large quantities of lactate by reducing pyruvate, a reaction that is catalyzed by lactate dehydrogenase (LDH) and coupled to the conversion of NADH to NAD^+^ (33). To test if enhanced shunting of pyruvate through SpxB might diminish lactate production, lactate levels were measured in the supernates of bacterial batch cultures prepared with TY medium supplemented with glucose or lactose as the carbohydrate source. The results showed reduced lactate levels, by about 40%, in cultures of the *manL* mutant grown on glucose, compared to the wild type grown on glucose (Fig. 1D); an outcome supporting that pyruvate was directed away from lactate generation. On the other hand, no significant differences in lactate accumulation were seen in cultures grown on lactose. It is likely that reduced homolactic fermentation by the *manL* mutant resulted in redirection from lactate production to acids with a higher p*Ka*, e.g., acetate, or other non-acidic end products, such as ethanol and acetoin, which may enhance the survival of *S. sanguinis* by increasing the intracellular pH and reducing the amount of damage caused by low environmental pH. However, this interpretation would not be entirely consistent with the modestly lower environmental pH achieved by the *manL* mutant when growing in BHI medium (Fig. 3C).

### Loss of *manL* alters central metabolism

To further characterize the role of the glucose-PTS in carbohydrate metabolism by *S. sanguini*s, RT-qPCR was performed to measure the mRNA levels of genes involved in central metabolism and related pathways in the *manL* mutant grown with glucose or lactose (Fig. 5 and Fig. S4). Consistent with the aforementioned phenotypes, transcriptional analysis indicated that the *manL* mutant, when growing on glucose, had reduced expression of the gene for lactate dehydrogenase (*ldh*). Also reduced was the expression of *pykF* gene, encoding for pyruvate kinase, which converts PEP into pyruvate with the concomitant generation of ATP. Reductions in mRNA levels of *pykF* and *ldh* are indicative of decreases in glycolytic rate and homolactic fermentation, respectively.

**Figure 5.**
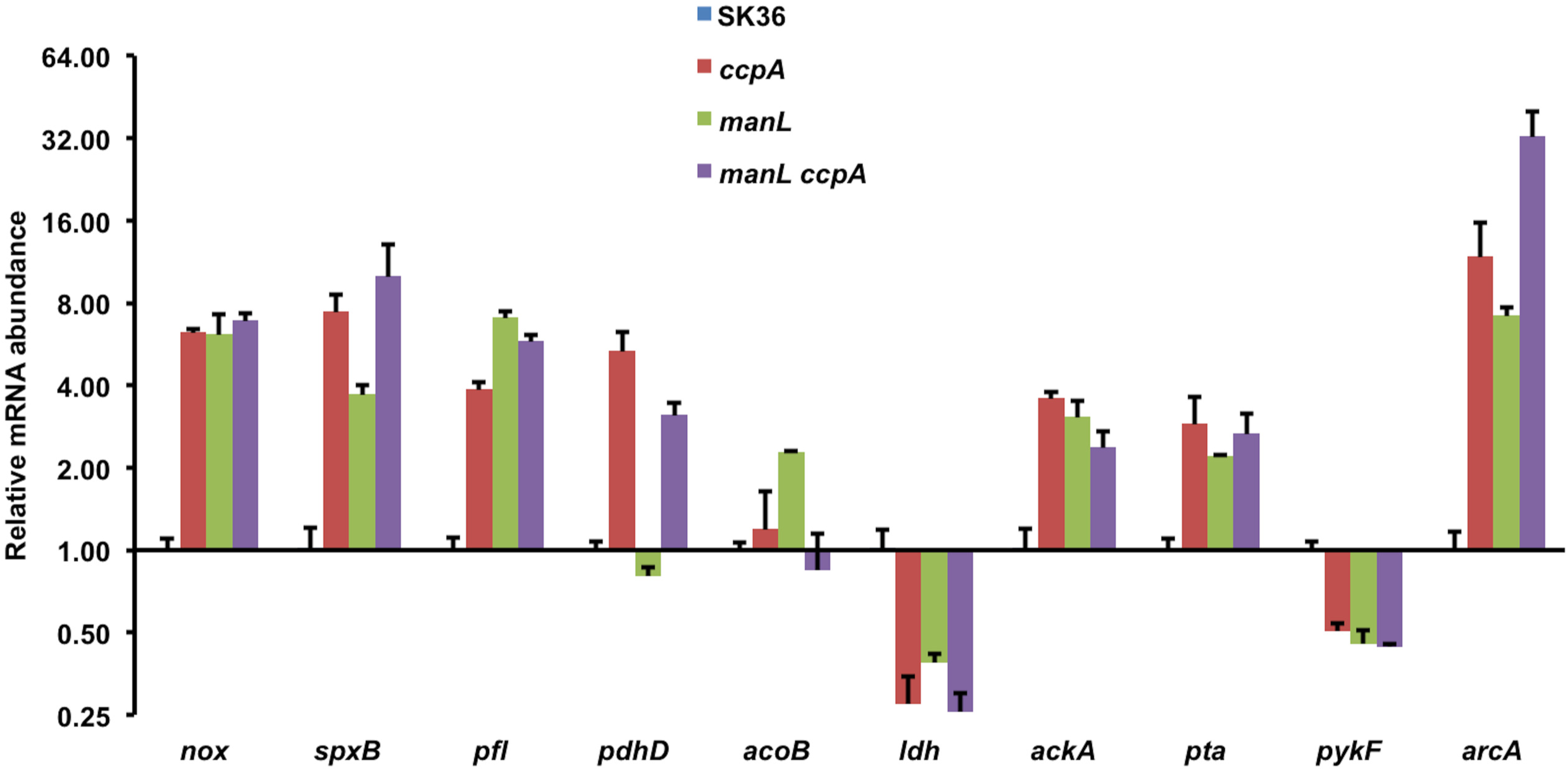
Measurements of relative mRNA levels of catabolic genes by RT-qPCR. Strains SK36, *ccpA*, *manL*, and *manL ccpA* were each cultured, in a TY medium containing 20 mM glucose, to exponential phase before harvested for RNA extraction. An internal control (*gyrA*) was used to measure the relative abundance of each transcript. The results for each gene are presented as the average and standard derivation (error bar) of three biological repeats.

Conversely, the *manL* mutant displayed enhanced expression by genes in oxidative pyruvate pathways that included *nox, spxB, pfl, acoB*, *pta,* and *ackA*. NADH oxidase (NOX) is required for oxidation of NADH into NAD^+^ in the presence of oxygen, resulting in formation of H_2_O. The levels of *pfl* mRNA, for a pyruvate-formate lyase that catalyzes the conversion of pyruvate into acetyl-CoA and formate, and *acoB* (SSA_1176), which belongs to the acetoin dehydrogenase operon, were also higher. As a neutral product of pyruvate catabolism, acetoin can be produced and released without influencing cytoplasmic or environmental pH, or it can be converted by the acetoin dehydrogenase complex into acetyl-CoA (34). Gene products of *pta* and *ackA* are required for further metabolism of acetyl-CoA, leading to production of acetate with concurrent generation of ATP. While SpxB is directly responsible for oxidation of pyruvate with production of H_2_O_2_, Nox was found to be required for optimal H_2_O_2_ release (35, 36). Enhanced expression of the genes for these enzymes substantiated earlier observations of increased H_2_O_2_ release by the *manL* mutant, as well as replacement of lactate by alternative end products, such as acetate. In each case, the *manL* mutant showed significant changes in gene expression when growing on glucose, but not on lactose, and the *manLComp* strain produced the mRNAs of interest at levels comparable to the wild type (Fig. S4).

### The *manL* mutant releases more pyruvate, but less eDNA

Catabolite control protein CcpA negatively regulates the expression of the pyruvate oxidase gene *spxB* and production of H_2_O_2_ in *S. sanguinis* strain SK36 (20). However, the *ccpA* mutant failed to show improved inhibition of *S. mutans* UA159 *in vitro*, a phenotype that was attributed to overproduction and release of pyruvate, an antioxidant that can scavenge H_2_O_2_ and thereby diminish its antagonism of *S. mutans* (21). Since deletion of the glucose-PTS impedes growth on glucose, it was reasoned that a *manL* mutant should experience a reduction in CCR, which could lead to production of elevated levels of H_2_O_2_ (as noted in Fig. 4) and perhaps even pyruvate. In contrast to a *ccpA* mutant, an increase in H_2_O_2_ secretion due to loss of *manL* was not evident when cells were grown on lactose, but the effects were especially evident when glucose, galactose, GlcN, or GlcNAc was the primary growth carbohydrate. In fact, the *ccpA* mutant failed to show any difference in H_2_O_2_ secretion from the wild type when grown on galactose or the amino sugars. It appeared that loss of *manL* affected H_2_O_2_ production more broadly than *ccpA* deficiency and the effects of mutations could manifest differently depending on the growth carbohydrate.

Significantly, the *ccpA* mutant failed to inhibit UA159 in our plate assays, as suggested previously (21), but the *manL* mutant inhibited UA159 moreso than the wild type on all carbohydrates tested except lactose (Fig. 4B). We also measured pyruvate levels in overnight TY-Glc batch cultures of the *manL* or *ccpA* mutant and the results (Fig. 6A) showed significantly more pyruvate in *manL* cultures (0.37 mM/OD_600_) than both the WT (0.07 mM/OD_600_) and the *ccpA* mutant (0.17 mM/OD_600_). No effect of either mutation was noted in cells grown on lactose. Monitoring of pyruvate release throughout the growth phases gave no indication that pyruvate was being actively reinternalized at any growth stage (Fig. 6BC); *S. mutans* actively reinternalizes pyruvate when it begins to enter stationary phase (37). The OD_600_ results (Fig. 6BC) from this experiment (also see Fig. S5A) revealed enhanced biomass of the *manL* mutant throughout the 24-h period. To explore the possibility that deletion of *manL* affected bacterial autolysis, extracellular DNA (eDNA) levels were measured in overnight TY-Glc cultures using SYTOX Green. Significantly lower eDNA levels were present in the cultures of the *manL* mutants than in cultures of the wild type (Fig. 6D), suggesting that the *manL* mutant lysed less than the wild type; when lysis of streptococci occurs, it is usually in stationary phase. Interestingly, a *manL ccpA* double mutant of SK36 produced pyruvate at levels comparable to the wild type (Fig. 6A). Furthermore, the *manL ccpA* double mutant showed a greater capacity to inhibit the growth of UA159 than the *ccpA* mutant (Fig. 4B). These results indicate that the ManL component of the glucose-PTS is capable of regulating central metabolism independently of CcpA.

**Figure 6.**
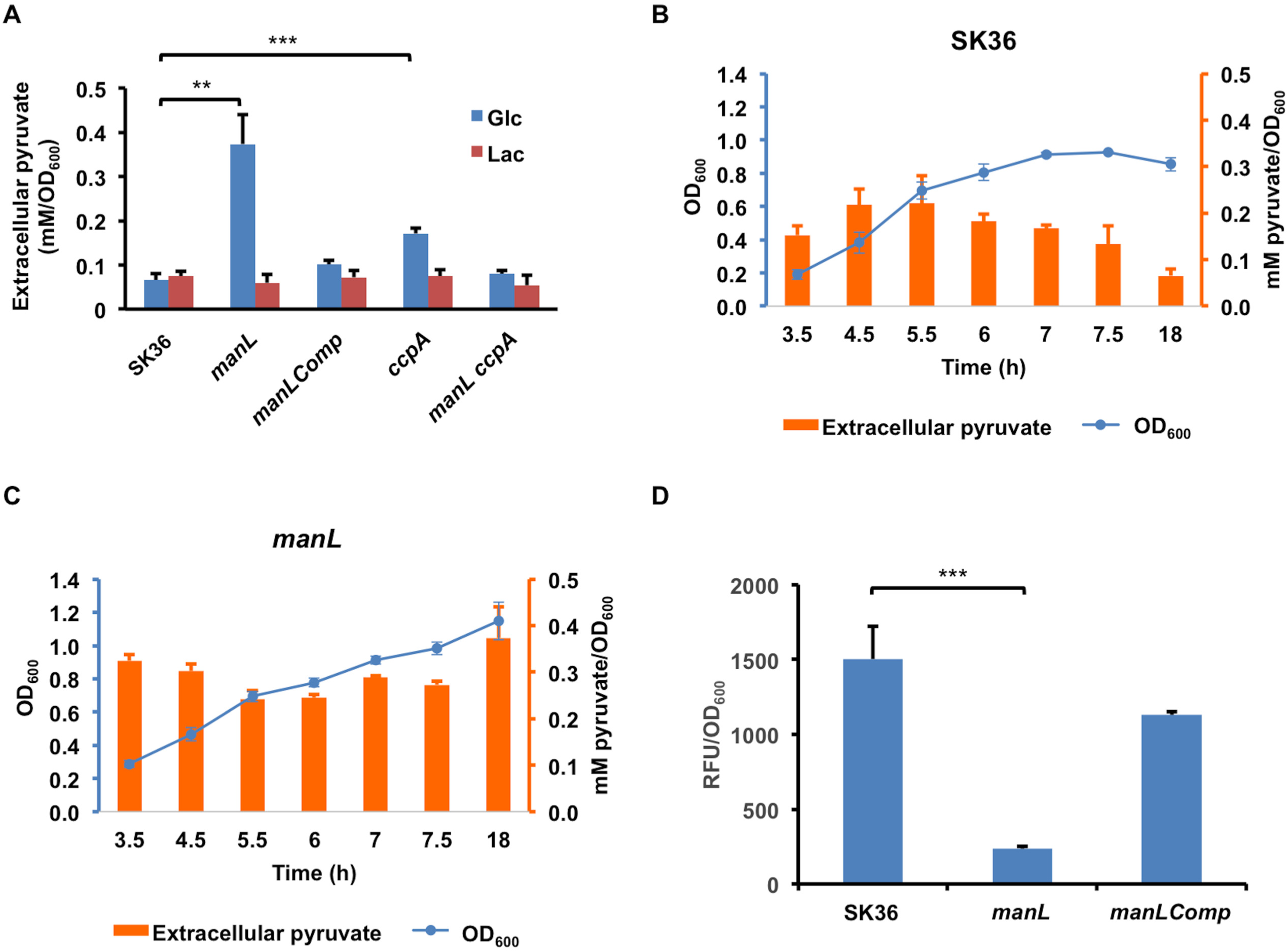
Deletion of *manL* affects release of pyruvate (A, B, C) and eDNA (D). Strains SK36, *manL*, *manLComp*, *ccpA*, and *manL ccpA* were cultured in TY medium supplemented with 20 mM of glucose (A, B, C, D) or 10 mM of lactose (A). (A) Supernates from stationary-phase (20 h) cultures of all 5 strains were used in an LDH-catalyzed reaction to measure the pyruvate levels under glucose and lactose conditions. Aliquots of SK36 (B) or the *manL* mutant (C) were taken at specified time points for measurements of OD_600_ and extracellular pyruvate. (D) Supernates from stationary cultures of SK36, *manL*, and *manLComp* were measured for DNA concentrations using a fluorescent dye. The relative fluorescence units (RFU) are in linear relationship with DNA concentrations within the experimental range. Each result shows the average of three biological repeats, with the error bars denoting standard derivation and asterisks the statistical significance according to Student’s *t* test (**, *P* <0.01; ***, *P* <0.001).

Transcriptional analysis was again performed in order to better test this hypothesis. The results of the RT-qPCR (Fig. 5) showed that most genes involved in pyruvate metabolism were regulated in similar fashions in the *manL* and *ccpA* mutants, except for pyruvate dehydrogenase (*pdh*) and acetoin dehydrogenase genes. Specifically, whereas the *ccpA* mutant produced about 5 times more *pdhD* mRNA than the wild type, the *manL* mutant showed no significant change in *pdhD* expression. The levels of *acoB* mRNA remained largely unchanged in the *ccpA* mutant but were increased > 2-fold in the *manL* mutant. Considering the significance of Pdh in metabolizing pyruvate under aerobic conditions, its enhanced expression in the *ccpA,* but not in the *manL* mutant, could provide an explanation for the higher levels of pyruvate being detected in the glucose cultures of the *manL* mutant. In addition, both the *ccpA* and the *manL* mutants produced significantly lower *ldh* mRNA than the wild type, consistent with the apparent redirection of pyruvate away from lactate production. It was not immediately clear why such high levels of extracellular pyruvate did not suppress H_2_O_2_-mediated antagonism of UA159 by the *manL* mutant (Fig. 4B). However, we hypothesized that accumulation of pyruvate itself could be a contributing factor to the enhanced fitness of the *manL* mutant in acidic environments and when the levels of H_2_O_2_ were elevated, especially in consideration that H_2_O_2_ can trigger autolysis and eDNA release in *S. sanguinis* (26, 38). Aside from being an antioxidant, metabolism of pyruvate via the oxidative pathways should yield additional ATP, which is beneficial to persistence, especially for maintaining pH homeostasis via the activity of F_1_F_o_- ATPase (39). Thus, the enhanced competitive fitness of a *manL* mutant of *S. sanguinis* may be attributable to a variety of factors that influence antagonism against *S. mutans*, as well as survival of *S. sanguinis* in the presence of *S. mutans* and its antagonistic products.

### Exogenous pyruvate benefits SK36

To test the effects of pyruvate on the fitness of *S. sanguinis*, SK36 was inoculated in BHI with or without 5 mM pyruvate and incubated overnight (≥ 20 h) before the cultures were diluted and enumerated for CFUs on agar plates. At the same time, the cultures were diluted (1:100) into TY-Glc with and without 5 mM pyruvate before growth was monitored. Strain UA159 of *S. mutans* was included for comparison, as it is known to be highly aciduric. In both assays, inclusion of pyruvate in overnight BHI cultures significantly enhanced the persistence of SK36, as assessed by CFU (*P* <0.05, Fig. 7A) and growth rate of subcultures in TY-Glc (Fig. 7B). Addition of pyruvate in TY-Glc also enhanced the growth rate of SK36 that was diluted from overnight cultures prepared without pyruvate (Fig. 7C). When the *manL* mutant of SK36 was cultured with added pyruvate, it too produced enhanced CFU counts, although its CFU counts were greater than the wild type with or without addition of pyruvate (Fig. 7A). By contrast, *S. mutans* UA159 showed no statistically significant change in either overnight viable counts or growth characteristics of TY-Glc cultures in response to the presence of 5 mM pyruvate. These results suggested that the presence of pyruvate in the environment could benefit the persistence of the commensal relative to *S. mutans*.

**Figure 7.**
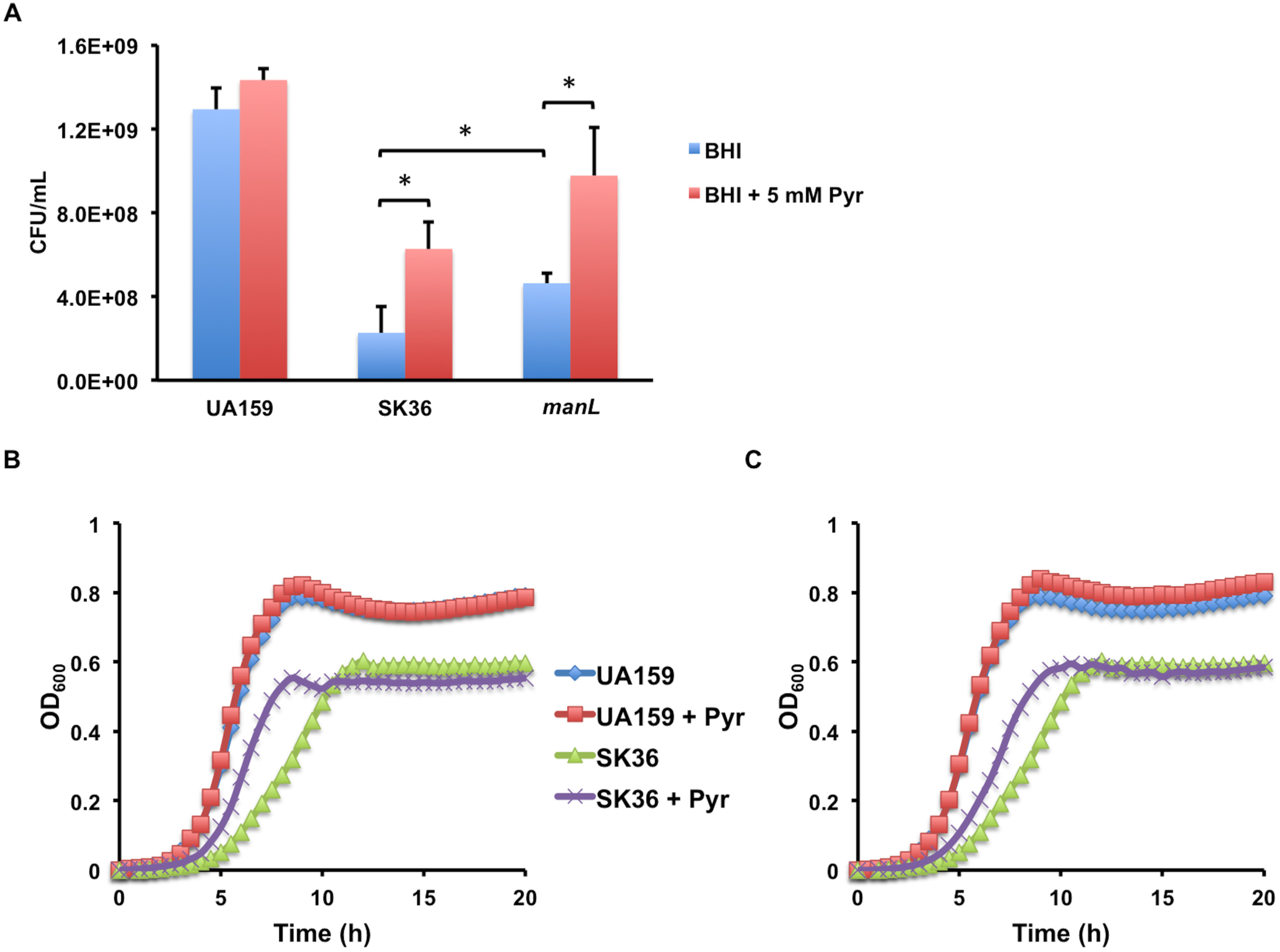
Exogenous pyruvate improves persistence of *S. sanguinis*. *S. mutans* UA159 and *S. sanguinis* strains SK36 and its isogenic *manL* mutant were each (n = 3) cultured overnight in BHI medium supplemented with or without 5 mM of pyruvate. The next day, cultures were diluted and plated for viable CFU counts (A) or for assessment of growth characteristics (B, C) by diluting into TY-Glc medium with or without 5 mM of pyruvate. Panel B shows the effects of pyruvate in overnight cultures of UA159 and SK36 by diluting them into TY-Glc without pyruvate, and panel C shows the effects of pyruvate added to fresh TY-Glc by diluting from overnight cultures prepared without pyruvate. Each of the columns and the growth curves represents the average of three biological replicates. Asterisks denote the statistical significance (*P* <0.05) obtained using a Student’s *t* test.

In light of the presence of *manL* mutations and the abundance of other SNPs (Table S1) detected in our two SK36 stocks, we obtained a third stock of SK36 from ATCC and constructed a *manL* deletion mutant for comparison. Studies of this new mutant in persistence, antagonism of UA159, release of H_2_O_2_ and pyruvate, as well as other growth characteristics showed effects highly similar to the *manL* mutants created using the other two strains of SK36 (Fig. S5 and data not shown). All three SK36 stocks also displayed highly similar phenotypes in these assays (Fig. S5). To our surprise, WGS indicated that the ATCC SK36 stock (here designated MMZ1922) carries nearly identical SNPs (totaling 116, Table S1) to those of the aforementioned two stocks, raising the possibility that the genome sequence of SK36 deposited at GenBank was derived from a strain that is significantly different from what is available from ATCC and what is being used by at least two oral streptococcus research groups. The basis for the differences between the sequence deposited in GenBank and that of the strains used here is not known, but could be accounted for by advances in sequencing technology since the original sequence was deposited. Meanwhile, an analysis using multi-locus sequence typing (Fig. S6) placed both the GenBank sequence and the assembled genome of the ATCC stock (MMZ1922) within the species of *S. sanguinis* as recently determined by sequencing and phylogenomic analysis of 25 low-passage clinical isolates of *S. sanguinis* (40).

### Detection of extracellular pyruvate in cultures of oral bacteria

Detection of spontaneous *manL* mutants in SK36 raised the possibility that isolates with similar genetic lesions could exist in nature, and those isolates would likely display similar phenotypes, including increased excretion of pyruvate. This hypothesis was tested using two cohorts of oral bacterial isolates made available from previous studies, including 63 commensal isolates from mostly caries-free donors (40) and 96 clinical isolates from both caries-free and caries-active subjects (41). When grown with TY-glucose or TY-lactose overnight in an aerobic atmosphere, a majority (∼90%) of these bacteria did not produce significant levels of pyruvate in their culture supernates. However, a total of 16 strains had significant levels of pyruvate (>0.15 mM/OD_600_) in their culture media 20 h post inoculation (Fig. S7A). These pyruvate-positive strains included multiple oral streptococcal species, from both caries-free and caries-active hosts. The issue of whether the original stocks of these isolates contained subpopulations of *manL* mutants was not evaluated because of the unrealistic scope of such an undertaking. Still, it must be considered that the results could reflect the sum of heterogeneous behaviors of genetically different subpopulations, the proportions of which may differ by isolate.

We next cultivated unstimulated whole saliva samples obtained from four healthy donors in a semi-defined medium containing various carbohydrates (42), including glucose, fructose, galactose, GlcN, GlcNAc, lactose, sucrose, or maltose, then measured pyruvate levels in the supernates. Out of four saliva samples, three produced significantly higher pyruvate levels on certain carbohydrates, with each showing a different profile in response to the primary carbohydrate in the medium (Fig. S7B). These latter results demonstrate that pyruvate can accumulate in complex populations of oral bacteria that more closely mimic the composition of the salivary microbiome in a carbohydrate-dependent manner.

## Discussion

In addition to acidogenicity, acid tolerance (aciduricity) is a key attribute of organisms that contribute to the initiation and progression of dental caries. The continued metabolic activities of acid-tolerant bacteria under acidic conditions shifts the ecological balance away from health-associated, acid-sensitive commensals, enhancing the virulence of the biofilms. While much attention has been paid to the aciduricity and acid tolerance response (ATR) by mutans streptococci, especially *S. mutans*, limited information is available regarding many of the abundant commensal streptococci. These bacteria make up the bulk of the dental microbiome and remain part of the biofilm even during significant caries events (43), and many are capable of fermenting carbohydrates as efficiently or superior to the etiological agents of caries (44, 45). Here we reported the identification of spontaneous mutants of *S. sanguinis* SK36 deficient, in whole or in part, in the ManL component of the glucose-PTS that showed improved fitness in stationary phase and acidic conditions. These phenotypic changes coincided with a shift of central carbon metabolism away from lactate generation in favor of pyruvate-oxidizing enzymes, resulting in increased secretion of H_2_O_2_ and pyruvate, and increased arginine deiminase activity. As such, the *manL* mutant of SK36 not only outcompeted its isogenic parent in stationary phase, but also showed significantly greater competitiveness against *S. mutans* on agar plates. As the conditions that likely contributed to the emergence of these mutants - growth in rich media containing high concentrations of glucose and the resultant low pH - are also expected in caries-conducive dental biofilms, our findings could have relevance in understanding the role of the glucose-PTS in regulating bacterial metabolism in ways that affect the overall aciduricity of dental biofilms. The findings also give rise to the hypothesis that there may be selection for mutants of commensal streptococci with defects in the ManL component of the PTS under cariogenic conditions that allow these variants to better survive, and perhaps even contribute to, cariogenic conditions.

Why is the *manL* mutant better at surviving in an acidic environment? When using fast-growing, exponential phase cells, pH drop and acid killing assays did not reveal the *manL* mutant to be more acid tolerant than the wild type. Nevertheless, it was clear that the mutant fared better under certain acidic conditions and was better able to maintain viability following a freeze-thaw cycle. We were also able to show that after prolonged incubation in the stationary phase, either on agar plates or in liquid cultures, the *manL* mutant had significantly greater viability than the wild type. It is possible that the most pronounced phenotype of the *manL* deletion occurs during the stationary phase, when the pH happens to be the lowest. For example, the *manL* mutant could have reduced propensity for autolysis, as our results have indicated (Fig. 6D), and it is known that extreme pH and/or high H_2_O_2_ levels can trigger cells to undergo autolysis (26, 38, 46). Based on qRT-PCR data, we hypothesize that a few mechanisms could be responsible for enhanced fitness of mutants lacking an intact ManLMN permease: i) altered metabolic end products, namely less lactate and more acetate, perhaps even change in acetoin levels, result in elevated intracellular pH with less damage to critical cellular components; ii) enhanced activities in pyruvate oxidative pathways provide more efficient production of ATP; iii) increased expression of the AD system allows cells to catabolize arginine, which benefits the bacterium both bioenergetically through creation of ATP and improved pH homeostasis; and iv) reduced LDH activity creates a surplus of pyruvate, which could scavenge H_2_O_2_ or provide a substrate to extend ATP production or be used for biogenesis pathways that enhance persistence under stressful conditions (e.g. membrane remodeling). Though not widespread in streptococci, *S. sanguinis* genome harbors putative genes required for gluconeogenesis which converts pyruvate into metabolic intermediates and precursors required for critical biogenesis pathways (16, 17). Indeed, pyruvate has been shown to resuscitate VBNC (viable but not culturable) *E. coli* cells by promoting macromolecular biosynthesis (47). Intracellular pyruvate could also potentially serve as a signal that alters gene regulation in favor of acid resistance (48). Further study on this subject, e.g. analysis at the systems level via proteomics or metabolomics, will be needed for confirmation of these theories.

Recently *S. mutans* was shown to release pyruvate as an overflow metabolite, which could then be reinternalized via dedicated transporters once excess glucose in the environment is depleted (37). *S. mutans* possesses two holin/antiholin-like proteins, LrgAB, that are responsible for transporting pyruvate at the start of stationary phase. Research has suggested the existence of additional transporters or mechanisms for pyruvate to impact the growth of *S. mutans* (49). The genome of *S. sanguinis* lacks homologues of either gene, but *S. sanguinis* and other *spxB*-encoding species do release pyruvate in varying amounts to the surroundings, which can decrease the damaging effects of H_2_O_2_ (21). Our measurements of exogenous pyruvate did not seem consistent with an highly active mechanism for reinternalization of pyruvate in SK36 (Fig. 6BC), yet addition of exogenous pyruvate improved bacterial persistence in overnight cultures (Fig. 7; Fig. S5B). Relative to the control, cultures with added pyruvate showed enhanced viability when sub-cultured. Meanwhile, pyruvate treatment failed to enhance the final OD_600_ or pH of overnight cultures, however it did reduce eDNA release by a modest amount (23%, *P* <0.05; Fig. S5ACD). Again, these results run counter to the existence of a dedicated pyruvate transporter for *S. sanguinis*, however they do not rule out that pyruvate uptake occurs nonspecifically. Alternatively, as H_2_O_2_ can diffuse freely through cell membranes, pyruvate could impact bacterial physiology simply by reacting with H_2_O_2_ in the environment.

How does the glucose-PTS regulate multiple metabolic pathways? Excess amounts of exogenous glucose can inhibit oxidative phosphorylation in favor of ethanol or lactate fermentation; a phenomenon termed the Crabtree effect in eukaryotic systems, e.g. yeast and tumor cells (50), or carbon catabolite repression (CCR) in facultatively anaerobic bacteria. The genetic mechanism responsible for this effect in most Gram-positive bacteria has been attributed to catabolite control protein CcpA, although recent research has pointed to other PTS-specific, CcpA-independent mechanisms, particularly in streptococci (19, 51–54). Our transcription analysis (Fig. 5) showed altered expression, due to loss of *manL,* in many central metabolic pathways, for which we envisioned three possible mechanisms. First, loss of *manL*, with the accompanying reduction in PTS activity could significantly impede PTS-mediated glucose influx, thereby reducing the steady state concentrations of the critical metabolic intermediates G-6-P and F-1,6-bP. In turn, the enzymatic activity of LDH, which is known to be allosterically activated by F-1,6-bP (10), also decreases. As LDH activity is critical to maintaining the NAD^+^/NADH balance, this change would likely affect the expression of multiple genes responsive to intracellular redox balance (36, 55), including *ldh* itself (56). This rationale could likewise be applied to cells growing in GlcN or GlcNAc, the catabolism of which yields F-6-P and subsequently F-1,6-bP. However, this would not apply in the case of galactose metabolism, as neither the tagatose pathway nor the Leloir pathway yields these intermediates. The second possibility involves CcpA-dependent regulation of gene expression. Since the ManLMN permease is the primary transporter of a number of monosaccharides, its activity is needed to sustain a certain level of energy intermediates, namely G-6-P, F-1,6-bP, and ATP, that are crucial to the function of the major catabolite regulator CcpA. A recent study of the CcpA regulon in *S. sanguinis*, performed using cells cultured in rich-medium, has revealed the various pathways and cellular functions that are controlled by CcpA (18). This list included *manL*, *pfl, spxB,* and several hundred other genes. The third scenario would be CcpA-independent. Our transcription analysis (Fig 5) indicated that *ldh*, *nox*, the *arc* operon, and the *pdh* operon are affected by the deletion of *manL* gene, however they were not identified as part of CcpA regulon in the aforementioned study (18); despite the fact that CcpA does control *arc* gene expression in the metabolically similar bacterium *S. gordonii* (8). Compared to the *ccpA* mutant, the *manL* mutant showed significantly more drastic effects on H_2_O_2_ production when growing on glucose, galactose, GlcN, GlcNAc, or a combination of glucose and galactose. Except for glucose or glucose and galactose, the *ccpA* mutant actually behaved much like the wild type (Fig. 4). Likewise, production of H_2_O_2_ by the *manL ccpA* double mutant matched that of the *ccpA* mutant in most cases, but was greater than that of the *ccpA* mutant on the amino sugars. Further, while a *nox* mutant showed a greatly reduced capacity to release H_2_O_2_, a *manL nox* double mutant produced more H_2_O_2_ than the *nox* mutant did under most conditions (Fig. 4). Although *spxB* is under the direct control of CcpA, our findings strongly support a CcpA-independent, PTS- specific pathway for regulating SpxB and related activities. A recent study on glucose-dependent regulation of *spxB* in *S. sanguinis* and *S. gordonii* identified significant distinctions between these two commensals in their response to availability of glucose (20), suggesting the presence of regulatory mechanisms beyond CcpA that could control pyruvate metabolism. We also previously reported that the glucose-PTS was capable of exerting catabolic control, independently of CcpA, over a fructanase gene (*fruA*) in *S. gordonii* (53).

In a caries-conducive dental plaque, rapid fermentation of dietary carbohydrates by lactic acid bacteria creates both detrimentally low pH and decreases the amount of carbohydrate available to commensals. These *in vivo* conditions do resemble that of a post exponential-phase batch culture whenever carbohydrate-rich media are used to grow streptococci. Such conditions could select for the *manL* mutations as the populations adapt to the stresses. In line with this reasoning, we have provided evidence supporting the existence of bacteria or mutants with decreased LDH activity and increased pyruvate oxidation and excretion within laboratory and clinical isolates, and shown variability in pyruvate production with a small set of *ex vivo* cultures grown from four different saliva donors. It is also understood that pyruvate exists in the oral cavity at varying levels, dependent upon the metabolic status and the functional makeup of the microbiome (57, 58). We recognize the fact that many of these isolates and multi-species cultures need further characterization in their metabolic capacities, as well as genetic composition. Nonetheless, the presence of these mutants in populations supports that there are advantages to the *manL* mutation, and perhaps other not-yet-identified mutations, under conditions such as those associated with caries development. Further, most of these bacteria are likely considered commensals, yet the impact of the physiological activities of these mutants on the dental microbiome remains unexplored. For example, *S. mutans* and other bacteria capable of actively transporting pyruvate could utilize exogenous pyruvate as a nutrient or for protection from oxidative stress. Previous studies in *S. mutans* show that the pyruvate dehydrogenase pathway is induced under starvation, contributing to aciduricity and long-term survival of the bacterium (59, 60). Further research into the role of PTS in pyruvate metabolism and its ecological impact to oral bacteria could reveal novel understandings in mechanisms that contribute to microbial homeostasis and oral health. Such knowledge of fundamental differences in the metabolism of carbohydrates by commensals and caries pathogens could be applied to promote health-associated biofilm communities.

## Materials and methods

### Bacterial strains and culture conditions

Three different stocks of *S. sanguinis* SK36 and their mutant derivatives, and *S. mutans* strain UA159 (Table 1) were maintained on BHI (Difco Laboratories, Detroit, MI) agar plates or Tryptone-yeast extract (TY, 3% Tryptone and 0.5% yeast extract) agar plates, each supplemented with 50 mM potassium phosphate, pH 7.2. Antibiotics including kanamycin (Km, 1 mg/mL), erythromycin (Em, 10 µg/mL), and spectinomycin (Sp, 1 mg/mL) were used in agar plates for the purpose of selecting for antibiotic-resistant transformants. BHI liquid medium was routinely used for preparation of batch starter cultures, which were then diluted into BHI, TY, or the chemically-defined medium FMC (61) modified to contain various carbohydrates at specified amounts. Cultures or the agar plates were incubated at 37°C in an aerobic environment with 5% CO_2_. Bacterial cultures were harvested at specified growth phases by centrifugation at 15,000 × *g* at 4°C for 10 min, or at room temperature for 2 min. The cells or the supernates were used immediately for biochemical reactions or stored at -80°C. For the purpose of studying growth characteristics, bacterial starter cultures were diluted into FMC or TY containing various carbohydrates and loaded onto a Bioscreen C system, where wells were overlaid with mineral oil, and cultures were maintained at 37°C.

Chromosomal DNA was extracted from bacterial cells using a Wizard Genomic DNA purification kit (Promega, Madison, WI), and submitted to MiGS (Microbial Genome Sequencing Center, Pittsburgh, PA; https://www.migscenter.com/) for WGS (Illumina) analysis and variant calling (Table S1). The coverage of these WGS genomes ranged between 40- to 50- fold. The high-throughput data of the genomic sequence of ATCC BAA-1455 (MMZ1922) were deposited in the Sequence Read Archive (SRA) and assigned an accession number PRJNA726918 (more information is available from the corresponding author upon request). To analyze the phylogenetic relationship of BAA-1455 with other streptococci, a multi-locus sequence typing (MLST) was performed using the genomes of *S. sanguinis* SK36 and *S. mutans* UA159 from GenBank, the ABySS assembly (14 Contigs) of MMZ1922 (62), along with the genomes of 25 low-passage *S. sanguinis* isolates and numerous clinical isolates of other streptococci obtained from our latest research (40).

### Construction of deletion mutants and complementing derivatives

An allelic exchange strategy (63) was modified to allow easy replacement of the target gene, e.g., *manL*, with multiple antibiotic markers (knockouts), and for the purpose of genetic complementation (*manLComp*), knocking-in of the wild-type gene (*manL*) in place of the antibiotic marker at the original site (26). Each recombinant event was facilitated by transformation of naturally competent bacterium using a linear DNA comprised of two homologous fragments, each at least 1 kbp in length, flanking an antibiotic marker (for knockouts) or a wild-type copy of *manL* sequence followed by a different antibiotic marker (for knock-ins). For the amplification of the upper flanking fragment, a 27-nucleotide sequence (sequence A) was added to the front of the regular reverse primer (Table S2, underlined in primer Ssa1918-2GA); and for the lower flanking fragment, a 30- nucleotide sequence (sequence B) was added to the front of the regular forward primer (underlined in primer Ssa1918-3GA). For PCR-amplification of antibiotic markers, including Km, Em and Sp, specific primers were designed for the integration of sequences A and B into the 5’- and 3’- ends, respectively, of each fragment (Table S2, labelled as “Marker for GA”). A mutator DNA for knockouts was created from two flanking DNA fragments and an antibiotic marker of choice, each at approximately 100 ng, using a 12-µL Gibson Assembly (GA) reaction (purchased from NEB or prepared in house) by incubation at 50°C for 1 h. Competent bacterial cells were induced by the use of a synthetic competence-stimulating peptide (CSP, by ICBR at the University of Florida) previously identified for *S. sanguinis* (64).

For complementation via knocking-in, a different “-2GA” primer (Ssa1918_manL_Comp-2GA) was designed and used together with the original forward primer (Ssa1918-1), so that the upper fragment now contained the wild-type gene in addition to the flanking sequence, ending with the same overlapping sequence A. This new fragment, the original lower fragment and an antibiotic marker of choice (different from the one replacing *manL*) were then ligated together via GA reaction to create a DNA to restore a wild-type *manL* operon. Each strain was confirmed by PCR using two outmost primers (with names ending in “-1” and “-4”; Table S2), followed by Sanger sequencing to ensure that no mutations were introduced into the target or flanking sequences. The primers used in sequencing, GA-Seq-5’ and GA-Seq-3’, were derived from the sequences A and B, respectively, each facing outward from the antibiotic marker, thus can be used for all mutants constructed the same way.

### PTS assay and pH drop

The capacity of the bacterium to transport carbohydrates and to lower environmental pH as a result of carbohydrate fermentation was assessed using the PTS assay (65, 66) or a pH drop assay, respectively, as previously described (39, 66).

### RNA extraction and RT-qPCR

Bacterial cultures (5 to10 mL) from mid-exponential phase (OD_600_ = 0.5-0.6) were harvested and treated with RNAprotect reagent (Qiagen, Germantown, MD), and the cell pellet, if not immediately processed, was stored at -20°C. Bacterial cells were resuspended in a lysis buffer (Qiagen), together with an equal volume of acidic phenol and similar volume of glass beads, and disrupted by beadbeating for 1 min. After 10 min of centrifugation at 15,000× *g* at room temperature, the clarified aqueous layer was removed and processed using an RNeasy mini kit (Qiagen) for extraction of total RNA. While loaded on membrane of the centrifugal column, the RNA sample was treated with RNase-free DNase I solution (Qiagen), twice, to remove genomic DNA contamination.

To synthesize cDNA, 0.5 µg of each RNA sample was used in a 10-µL reverse transcription reaction set up using the iScript Select cDNA synthesis kit (Bio-Rad), together with gene-specific reverse primers (Table S2) used at 200 nM each. Primer for the housekeeping gene *gyrA* was used as an internal control in all cases (67). After a 10-fold dilution with water, the cDNA was used as a template in a quantitative PCR (qPCR) reaction prepared using a SsoAdvanced Universal SYBR Green Supermix and cycled on a CFX96 Real-Time PCR Detection System (Bio-Rad), following the supplier’s instructions. Each strain was represented by three biological replicates and each cDNA sample was assayed at least twice in the qPCR reaction. The relative abundance of each mRNA was calculated against the housekeeping gene using a ΔΔCq method (68).

### H_2_O_2_ measurement and plate-based competition assay

The relative capacity of each strain to produce H_2_O_2_ and to compete against *S. mutans* was studied by following previously published protocols with minor changes. H_2_O_2_ production was assessed using an indicator agar plate on the basis of Prussian blue formation (21, 28). Briefly, a tryptone (3%)-yeast extract (0.5%) agar (1.5%) base was prepared with the addition of FeCl_3_.6H_2_O (0.1%) and potassium hexacyanoferrate (III) (0.1%). After autoclaving, glucose or other carbohydrates were added at the specified amounts before pouring plates. Each strain was cultivated overnight in BHI medium and dropped onto the agar surface, then incubated for >20 h to allow bacterial growth and development of Prussian blue precipitation. Each strain was tested at least three times, with 2 plates each time.

Plate-based inhibition assays (42) were carried out to test the interactions between *S. sanguinis* and *S. mutans* UA159 on various carbohydrate sources using TY-agar as the base medium. Overnight cultures of *S. sanguinis* strains were dropped onto the agar first, incubated for 24 h at 37°C in a 5% CO_2_ aerobic incubator, followed by spotting of *S. mutans* UA159 in close proximity to the *S. sanguinis* colony. Plates were incubated for another day before photographing. Each interaction was tested at least three times.

### Lactate and pyruvate measurement

Bacterial cultures were prepared by diluting overnight BHI cultures at 1:50 into TY medium supplemented with 20 mM glucose, then incubated at 37°C in an aerobic incubator maintained with 5% CO_2_. At the specified time or phase of growth, aliquots of bacterial cultures were taken for optical density measurement (OD_600_), or spun down using a tabletop centrifuge (14,000× *g*, 2 min), with supernates being removed for assays or stored at -20°C. Lactate levels in the culture supernates were measured using a lactate assay kit (LSBio, Seattle, WA), following the protocols provided by the supplier.

To measure pyruvate levels in the same cultures, we adopted an LDH-catalyzed reaction that coupled the reduction of pyruvate with the oxidation of NADH with monitoring of the optical density at 340 nm (OD_340_). The assay was performed by mixing 10 µL sample and 90 µL of enzyme solution, which included 10 Units/mL of lactate dehydrogenase (Sigma) and 100 µM NADH in a 100 mM sodium-potassium phosphate buffer (pH 7.2) supplemented with 5 mM MgCl_2_. The reaction was incubated at room temperature for 10 min before spectrometry, with the light source set at UV range. To rule out the influence to the assay by background NADH-oxidizing activities in bacterial cultures, a control without addition of LDH was included for each sample. A sodium pyruvate standard in the range of 0.05 to 0.6 mM was prepared freshly in TY base medium. The final measurements of pyruvate concentration were normalized against the optical density (OD_600_) of each culture. An earlier approach (69) to quantifying pyruvate using a commercial kit was not used due to the presence in samples of H_2_O_2_ that interferes with the reaction. Collection of saliva was carried out according to an established procedure (IRB201500497 at University of Florida) described elsewhere (42).

## Acknowledgements

This work was supported by DE12236 from the United States National Institute of Dental and Craniofacial Research. We thank Dr. Jacqueline Abranches for providing a subset of the clinical isolates of oral streptococci used in this study.

